# Biochemical adaptations of the retina and retinal pigment epithelium support a metabolic ecosystem in the vertebrate eye

**DOI:** 10.1101/143347

**Authors:** Mark A. Kanow, Michelle M. Giarmarco, Connor Jankowski, Kristine Tsantilas, Abbi L. Engel, Jianhai Du, Jonathan D. Linton, Christopher C. Farnsworth, Stephanie R. Sloat, Ken J. Lindsay, Edward D. Parker, Susan E. Brockerhoff, Martin Sadilek, Jennifer R. Chao, James B. Hurley

## Abstract

Here we report multiple lines of evidence for a comprehensive model for retinal energy metabolism. Metabolic flux, locations of key enzymes and our finding that glucose enters the neural retina almost entirely through photoreceptors support a conceptually new model for retinal metabolism. In this model, glucose from the choroidal blood supply passes through the retinal pigment epithelium to the retina where photoreceptors convert it to lactate. Photoreceptors then export the lactate as fuel for the retinal pigment epithelium and for neighboring Müller glial cells. A key feature of this model is that aerobic glycolysis in photoreceptors produces lactate to suppress glycolysis in the neighboring retinal pigment epithelium. That enhances the flow of glucose to the retina by minimizing consumption of glucose within the retinal pigment epithelium. This framework for metabolic relationships in retina provides new insights into the underlying causes of retinal disease, age-related vision loss and metabolism-based therapies.

## INTRODUCTION

Mutations in any of more than140 genes can cause photoreceptors in a vertebrate retina to degenerate (1). The relationship between photoreceptor degeneration and the diversity of biochemical functions and expression patterns of those genes is enigmatic. The diversity suggests that the consequences of loss or gain of function of these genes may converge onto a few essential metabolic processes (2, 3). Much has been gained by studying specific functions of the genes. Based on those studies therapeutic strategies for specific genetic deficiencies are being developed (4). Nevertheless, a more general approach to understanding what photoreceptors need to survive could lead to more broadly applicable therapeutic strategies. With that in mind, we have been investigating the fundamental nature of energy metabolism in the retina and the retinal pigment epithelium (RPE) (5–11).

Glucose that fuels the outer retina comes from the choroidal blood. Before it can reach the retina, however, the glucose first must traverse the RPE. The RPE is a monolayer of polarized cells between the choroid and retina that functions as a blood-retina barrier. The cells in the RPE are bound together by tight junctions and they express specific transporter proteins on their basolateral and apical surfaces (12). Glucose from the choroid passes through transporters on the basolateral surface, and then wends its way through the cytoplasm of the RPE cell. If metabolic enzymes within the RPE cell do not consume it, glucose moves down a concentration gradient toward the opposite side of the RPE cell where it exits to the retina through transporters on the apical surface of the RPE.

When the glucose reaches the retina most of it is consumed by glycolysis and converted to lactate. Retinas and tumors were the two tissues first identified in the 1920’s by Warburg (13) as relying mostly on “aerobic glycolysis”. This type of metabolism can release massive amounts of lactate from a cell even when O_2_ is available. Evidence indicates that photoreceptors in the outer retina are the site of aerobic glycolysis (8, 13–17). The importance of aerobic glycolysis for survival and function of photoreceptors has not been established, but it is thought to enhance anabolic activity within the photoreceptor (3, 18–20).

In RPE cells energy metabolism is strikingly different than in the retina. In particular, RPE cells are specialized for a type of energy metabolism called reductive carboxylation (9) that aids in redox homeostasis. In general, RPE cells rely more on their mitochondria.

Recent reports described genetic manipulations that explored the effects of qualitatively altering energy metabolism either in photoreceptors or in RPE cells *in vivo*. Glycolysis in rods was enhanced in one study by blocking expression of SIRT6 (3). Another study enhanced glycolysis in rods by activating mTORC1 (21). Those studies showed that making photoreceptors more glycolytic makes them more robust. Both strategies slowed degeneration of rods caused by mutations associated with retinitis pigmentosa (3, 21). However, making RPE cells more glycolytic *in vivo* has the opposite effect; it causes neighboring photoreceptors to degenerate. When glycolysis in the RPE was enhanced by knocking out VHL (22) or by knocking out an essential mitochondrial transcription factor in RPE cells *in vivo* (23) the neighboring photoreceptors died.

The findings of those *in vivo* studies appear puzzling and seemingly contradictory when considered only from a cell autonomous perspective. Why does enhancing glycolysis help some cells and endanger others? Here we propose that those findings make more sense when interpreted in the context of metabolic relationships between the retina and the RPE. We describe evidence that the retina and RPE function as a metabolic ecosystem. We show that photoreceptors are the main point of entry for glucose into the retina. The photoreceptors convert glucose to lactate, which then serves as a fuel for other cells in the retina. We show here that lactate also can enhance the ability of RPE cells to pass glucose from the blood to the retina. The model that we propose based on these findings predicts that each cell in the retina and RPE contributes an essential metabolic function that promotes survival of the complete retina-RPE ecosystem.

## RESULTS

### Photoreceptors express a glucose transporter

Uptake of glucose into cells requires a protein that can transport glucose. We used immunoblotting of mouse tissues to evaluate expression of glucose transporters (**Fig. 1A**) and confirmed previous findings that GLUT1 is the primary glucose transporter isoform in retina (24) and RPE (25). The protein immunoreactive with the GLUT1 antibody was confirmed to be membrane associated (**Fig. 1B**). GLUT3 was detected only in brain. GLUT4 was detected in heart and muscle as expected, but not in the retina.

**Fig. 1.**
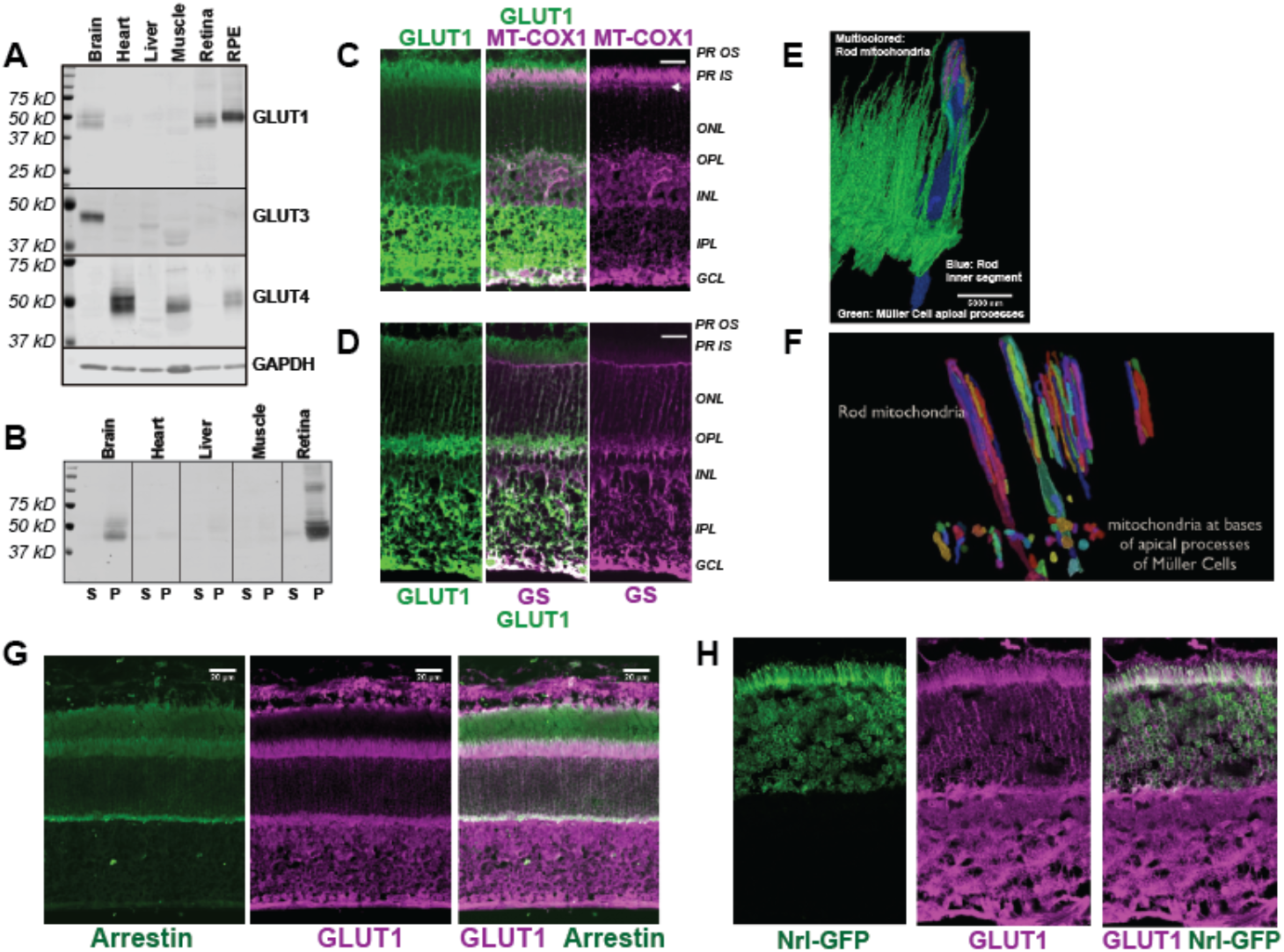
Distribution of GLUT1 in retina. **A.** Immunoblot analysis of mouse tissue homogenates confirms that GLUT-1 is the primary glucose transporter in mouse retina and RPE. 1 μg total homogenate protein was loaded in each lane. No antibodies that we could validate were available for GLUT2. However, the Human Protein Atlas reports no expression of GLUT2 in retina (53). **B**. Evidence that the protein immunoreactive with the GLUT-1 antibody is membrane associated. Homogenates were centrifuged and equivalent percentages of total supernatant (S) and total pellet (P) were probed with the GLUT-1 antibody. **C.** GLUT1 in mouse retina. Rods inner segments are identified by the unique morphology of their mitochondria labeled with mitochondrial cytochrome oxidase 1 antibody (MT-COX1). White arrowhead indicates the layer of Müller cell mitochondria. **D**. Müller cells are identified by glutamine synthetase (GS) immunoreactivity. Scale bars represent 20 μm. **E**. Serial Block Face Scanning Electron Microscopy. The inner segment of one rod cell is shown in blue with its mitochondria shown as multi-colored. The green structures are apical processes of MGCs. **F**. Differences in location and morphology between rod mitochondria and MGC mitochondria in mouse retina. For clarity not all of the mitochondria are shown. MGC mitochondria are located just below the outer limiting membrane. **G**. The left panel shows labeling of rods in a partially light-adapted mouse retina with a rod arrestin antibody. The middle panel shows labeling with a GLUT1 antibody and the right panel shows the merge **H.** The left panel shows expression of GFP from the rod-specific Nrl promoter. The middle panel shows labeling with a GLUT1 antibody and the right side shows the merge. PR OS, photoreceptor outer segment; PR IS photoreceptor inner segment; ONL, outer nuclear layer; OPL, outer plexiform layer; INL inner nuclear layer; IPL inner plexiform layer; GCL, ganglion cell layer.

Immunohistochemistry shows that GLUT1 immunoreactivity overlaps with cytochrome oxidase subunit 1 (MT-CO1) (**Fig. 1C**), which identifies rod inner segments by the unique elongated shape of their mitochondria (**Fig 1E)**. These mitochondria extend beyond the ends of the Müller glial cell (MGC) apical processes (**Fig 1E)**. There are no mitochondria within these fine MGC apical processes. Instead, small spherical-shaped mitochondria line up within the MGCs along the outer limiting membrane just beneath the apical processes (**Fig. 1F and arrowheads in Fig. 1C**). MGCs extend from the outer limiting membrane to the ganglion cell side of the retina. Most of the GLUT1 immunoreactivity in MGCs is in the inner retina (**Fig. 1D).** GLUT1 immunoreactivity also overlaps with a marker specific for rod photoreceptors, rod arrestin (**Fig. 1G**) and it overlaps with GFP expressed from the rod-specific Nrl promoter (**Fig. 1H**). Taken altogether, the distribution of GLUT1 immunoreactivity supports the idea that photoreceptors can take up glucose released from the apical side of the RPE.

### Dietary glucose enters the retina primarily through photoreceptors

Next we asked which cells in the retina take up glucose in the context of an eye within a living animal. We used oral gavage to introduce a fluorescent derivative of 2-deoxy glucose (2-NBDG) (26) into stomachs of mice. We harvested the retinas 20 or 60 minutes after gavage, mounted them on filter paper and cut slices for imaging by confocal microscopy (27). **Fig. 2A** shows that 2-NBDG fluorescence is strongest in the photoreceptor layer, suggesting that photoreceptors are the first cells in the retina to take up glucose from the blood. Surprisingly, 2-NBDG fluorescence is stronger in the outer retina than in the inner retina even though mouse inner retinas are vascularized. We noted that 2-NBDG fluorescence does not overlap with Müller Glial Cells (MGC’s), which were labeled in these experiments by transgenic expression of tdTomato (28), though in rare instances there was overlap at a MGC end foot.. These results are summarized and quantified in **Fig. 2C**. They show that glucose that reaches the outer retina is taken up primarily by photoreceptors and not MGC’s.

**Fig. 2.**
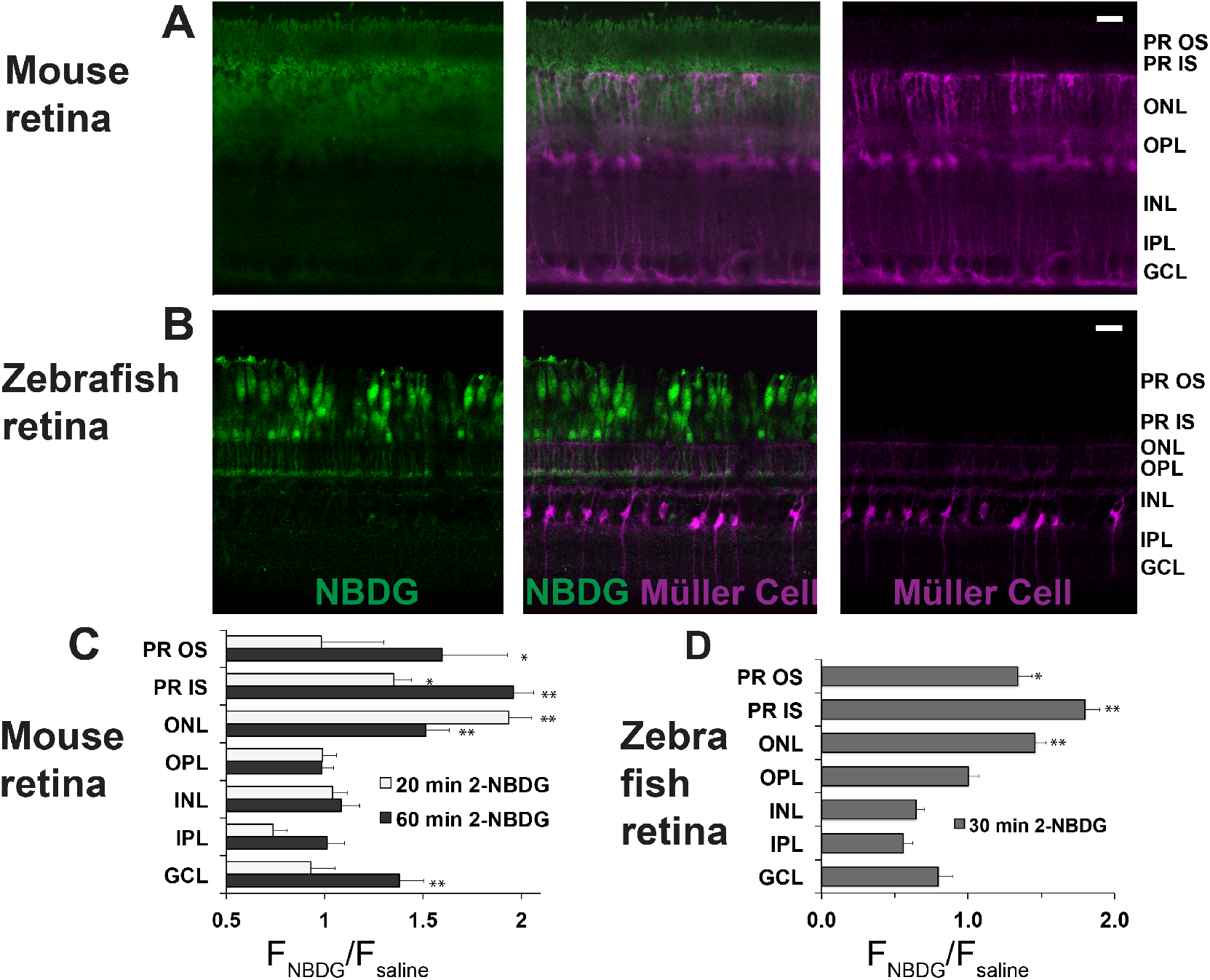
Fluorescent glucose (2-NBDG) accumulates in photoreceptors after oral gavage. **A.** 2-NBDG (green) accumulation in a mouse retina 20 min. after oral gavage. MGCs are identified by tdTomato in cells in which the Rlbp1 promoter is active. **B**. 2-NBDG accumulation in a zebrafish retina 30 min after oral gavage. MGCs are identified by tdTomato expressed from the GFAP promoter. Labels on the right of panels A and B represent approximate positions of the retinal layers, **C.** Quantification of 2-NBDG fluorescence from mouse retinas (n = 5 animals, 17 slices for 20 min 2-NBDG; 3 animals, 8 slices for 1 h 2-NBDG; 3 animals, 8 slices for saline). F_NBDG_/F_saline_ compares fluorescence from retinas of mice gavaged with 2-NBDG vs. with saline. Error bars report SEM. **D.** Quantification of 2-NBDG fluorescence from zebrafish retinas (3 animals, 8 slices for 30 min 2-NBDG; 2 animals, 3 slices for saline). PR OS, photoreceptor outer segments; PR IS, photoreceptor inner segments; ONL, outer nuclear layer; OPL, outer plexiform layer; INL, inner nuclear layer; IPL, inner plexiform layer; GCL, ganglion cell layer. Scale bars represent 20 μm.* indicates p < 0.05 and ** indicates p < 0.01 for the comparison of F_NBDG_ to F_saline._

The images in **Fig. 2A** were made from live, unfixed mouse retinas. Most photoreceptors in mouse retinas are rods. It is difficult in these images to resolve whether cones also import 2-NBDG. To address this we also introduced 2-NBDG by oral gavage into adult zebrafish, whose retinas are more enriched with cones (29). **Fig. 2B** shows that cones become intensely fluorescent 30 minutes after gavage. As in mouse retinas, there was no indication of glucose uptake into MGCs, which in these retinas were marked with tdTomato expressed from a GFAP promoter (30). **Fig. 2D** reports quantification and summarizes the zebrafish retina results.

### Carbons from glucose are metabolized in RPE cells more slowly than in retina

Previous studies showed that most of the glucose taken up into a retina is used to make lactic acid (8, 13–17). Within the eye of a living animal, glucose from the choroidal blood first must pass through the monolayer of RPE cells before it can reach the retina. We hypothesized that the energy metabolism of RPE cells might be adapted to minimize consumption of glucose in order to maximize the amount of glucose that can pass through the RPE to reach the retina.

To examine glucose metabolism in RPE versus in retina, we compared two preparations, mouse retina (mRetina) and cultured human fetal RPE cells (hfRPE). The mouse retinas were freshly dissected from mouse eyes. The hfRPE cells were grown 4-6 weeks in culture to form a monolayer with tight junctions and a trans-epithelial resistance similar to native human RPE (≥200 Ω• cm^2^). This hfRPE preparation has been widely used to study RPE metabolism and to model RPE-related diseases such as age-related macular degeneration due to its similarity to native RPE cells (31–35). We added ^13^C labeled glucose to both preparations and then used mass spectrometry (7) to compare incorporation of ^13^C into glycolytic and TCA cycle intermediates. For these experiments we used 1,2 ^13^C glucose because the pattern of ^13^C labeling from this isotopomer can be used to distinguish metabolites generated by glycolysis from metabolites generated by the pentose phosphate pathway (36). Metabolites with one ^13^C (“m1”) are generated from glucose that flows through the oxidative reactions of the pentose phosphate pathway whereas metabolites with two ^13^C (“m2”) are produced when glucose enters glycolysis directly. In a previous report we used this labeling method to show that < 2% of the metabolic flux from glucose goes through the pentose phosphate pathway in both mRetina and in hfRPE (9).

**Fig. 3A** shows the total pmoles per μg protein of several metabolites in mRetina and in hfRPE. There are several striking differences. Lactate and succinate are more abundant in mRetina than in hfRPE, whereas citrate and *α*-ketoglutarate are more abundant in hfRPE than in mRetina. **Fig. 3B** shows the time course of incorporation of ^13^C into several key metabolites. The initial rate at which ^13^C from glucose incorporates into intracellular lactate in mRetina is at least 8 times faster than in hfRPE. We also noted that the citrate and *α*-ketoglutarate pools are larger and fill more gradually in hfRPE cells than in retina indicating a large oxidative metabolic capacity of RPE mitochondria.

**Fig. 3.**
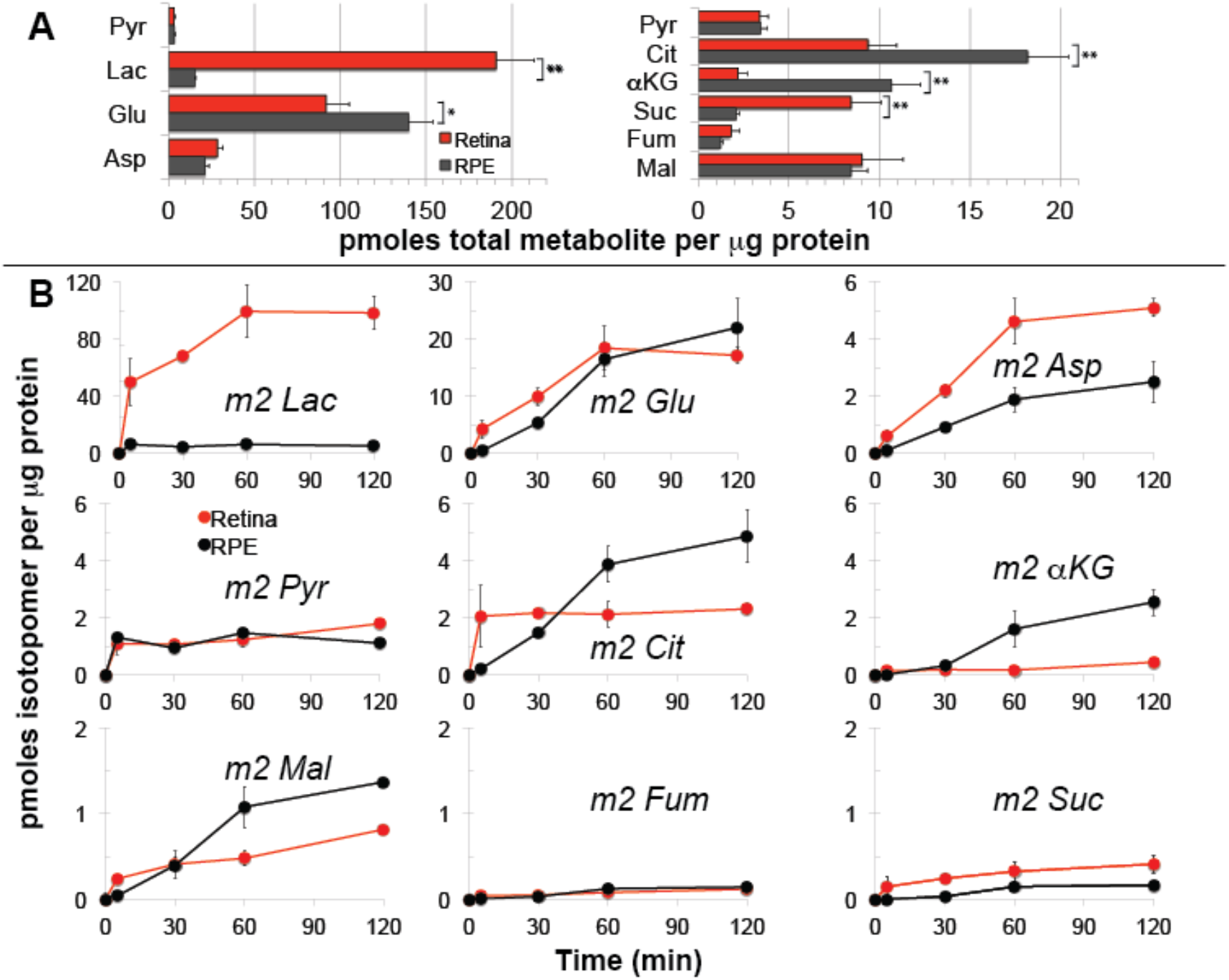
Differences in metabolic flux in retina and RPE. **A**. Total metabolite levels (pmoles per μg protein) in mRetina (red) and hfRPE (black). (n = 11) Note the different scales for the left and right panels. **B**. Incorporation of ^13^C from [1,2] ^13^C glucose into metabolites in mRetina and hfRPE cells (pmoles per μg protein). Each of the isotopomers shown is derived from glucose metabolized by glycolysis. Note the different scales for the top, middle and bottom panels. (n = 3 for each time point; error bars represent standard deviation).

These findings support our hypothesis that a slow rate of conversion of glucose to lactate in the RPE minimizes consumption of glucose so that more glucose can reach the retina. In a previous study we showed that RPE cells further minimize their use of glucose by using an alternative pathway known as reductive carboxylation to make NADPH (9). We propose that limited production of lactate and the use of reductive carboxylation to make NADPH are adaptations of RPE cells that minimize their consumption of glucose so that more glucose reaches the retina.

### RPE cells can use lactate as a fuel

The retina converts most of the glucose it consumes into lactate so we considered the possibility that RPE cells can use retinal lactate as an alternative fuel source to compensate for their minimal consumption of glucose, or even to further suppress glycolysis. We incubated hfRPE cells either with U-^13^C glucose or with U-^13^C lactate for 5 or 10 minutes and quantified incorporation of ^13^C into glycolytic and TCA cycle metabolites. **Fig. 4A** shows that ^13^C is incorporated rapidly into the pyruvate pool from both glucose and lactate. However, in the citrate pools, ^13^C from lactate accumulates at least 20 times faster than ^13^C from glucose. We also noted that substantial amounts of m3 malate are formed. This shows that carboxylation of pyruvate is a major metabolic pathway in hfRPE cells. **Fig. 4B** reports the time course of incorporation of ^13^C from lactate into TCA cycle intermediates in hfRPE cells. Carboxylation of pyruvate occurs at about half the rate of decarboxylation of pyruvate.

**Fig. 4.**
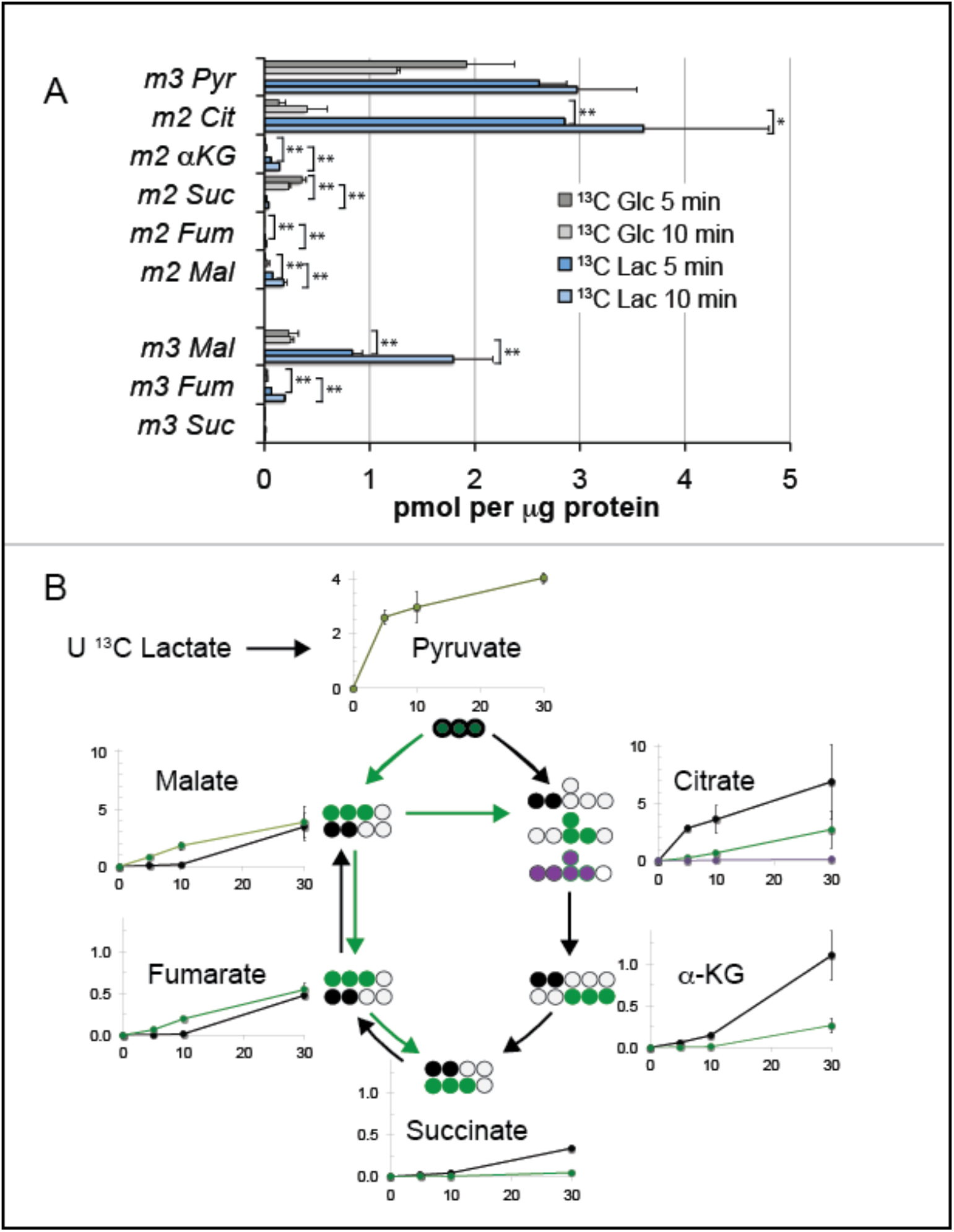
Incorporation of ^13^C from lactate into metabolic intermediates in hfRPE cells. **A**. Comparison of initial rates of labeling (at 5 and 10 minutes after introduction of labeled fuel) from U-C glucose vs. from U-C lactate. Citrate and malate take up label faster from lactate than from glucose. **B**. Time courses of incorporation of C from U-C lactate into hfRPE metabolites accompanied by schematic illustrations of the labeled species in the context of the TCA cycle. (n = 2-3 for each time point; error bars represent range or standard deviation).

### Lactate can suppress consumption of glucose by RPE cells

The results in **Fig. 3** and **Fig. 4** show that glucose is consumed more slowly by RPE cells than by retina. They also show that RPE cells efficiently use lactate for fuel as an alternative to using glucose. Based on these findings we next asked whether lactate can suppress consumption of glucose by RPE cells. We hypothesized (**Fig. 5A**) that lactate dehydrogenase (LDH) in RPE cells depletes cytosolic NAD^+^ when it oxidizes lactate to pyruvate. Since NAD^+^ is required for glycolysis, depletion of NAD^+^ by lactate and LDH would suppress glycolysis so that RPE cells consume less glucose.

**Fig. 5.**
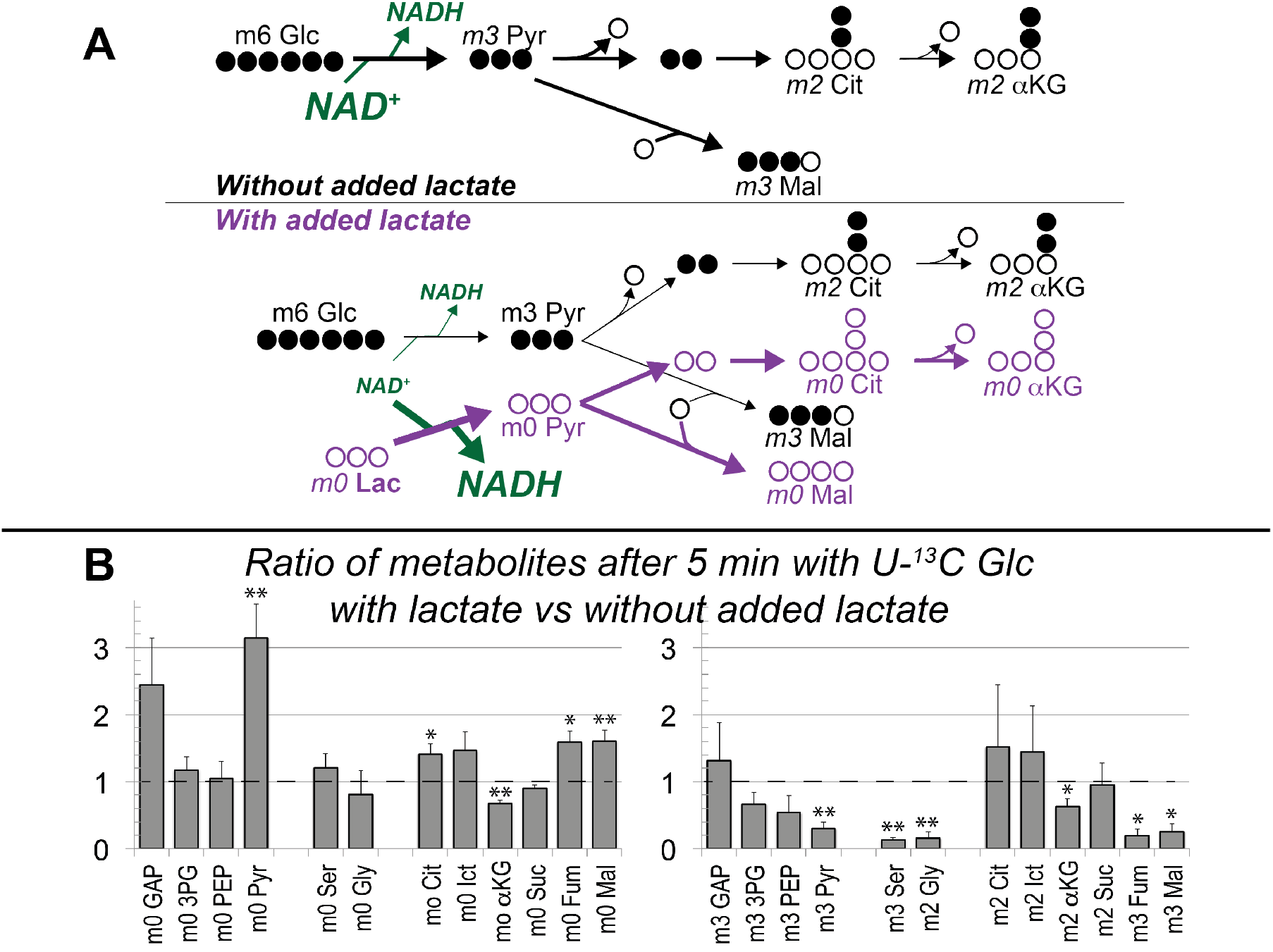
Lactate suppresses oxidation of glucose by hfRPE cells. **A**. Schematic prediction of how U-^13^C Glc (“m6 Glc”) would be metabolized without lactate (top) vs. with lactate (bottom). We hypothesized that lactate would suppress glycolysis of m6 Glc by depleting NAD^+^. The model also predicts that unlabeled (m0) pyruvate and TCA cycle intermediates become more abundant. **B**. Experimental results showing that lactate substantially increases the total amounts of unlabeled (m0) GAP, pyruvate, citrate, isocitrate, fumarate and malate (left panel) in hfRPE cells. The right panel shows that 20 mM lactate suppresses the incorporation of ^13^C from 5 mM ^13^C Glc into glycolytic and TCA intermediate. (n = 3; error bars represent SEM, * indicates p < 0.05 and ** indicates p < 0.01 for the comparison of with vs. without added unlabeled lactate.

We incubated hfRPE cell monolayers with 5 mM U-^13^C glucose either in the absence or presence of 20 mM unlabeled lactate. We then harvested the cells and used GC-MS to determine if lactate suppresses incorporation of ^13^C from glucose into glycolytic and TCA cycle intermediates.

The results of this experiment (**Fig. 5B**) show that unlabeled lactate substantially increases the amounts of unlabeled pyruvate, citrate, isocitrate, fumarate and malate (left panel of **Fig. 5B**). This is consistent with the results in **Fig. 4** showing that carbons from lactate are incorporated rapidly into TCA cycle metabolites through both carboxylation and decarboxylation of pyruvate.

Addition of lactate also causes accumulation of GAP, the triose phosphate immediately upstream of the glyceraldehyde-3-phosphate dehydrogenase (GAPDH) reaction, that requires NAD^+^. Consistent with this, added lactate diminishes the amount of ^13^C from glucose incorporated into intermediates downstream of the GAPDH reaction (right side of **Fig. 5B**). Lactate does not decrease incorporation of ^13^C from glucose into m2 citrate and m2 isocitrate, most likely because of anapleurotic supplementation of unlabeled TCA cycle intermediates through pyruvate carboxylation enhanced TCA cycle activity (see left side of **Fig. 5B**). Overall, lactate substantially inhibits glycolysis in hfRPE cells.

### Lactate can enhance transport of glucose across a monolayer of hfRPE cells

Next we asked whether lactate from photoreceptors can enhance the flow of glucose across the RPE. We reasoned that suppression of glycolysis by lactate minimizes consumption of glucose, so more glucose would cross from the basolateral to the apical side of the RPE.

To test this idea, we measured the influence of lactate on transport of glucose across a monolayer of hfRPE cells. We grew hfRPE cells on transwell filters to confluence with a transepithelial resistance ≥200 Ω• cm^2^. We then added ^13^C glucose to the basolateral side, where the RPE normally would face the choroid in an eye, and used mass spectrometry of the apical medium to quantify transport of ^13^C glucose to the apical side, where the RPE normally would face the lactate-rich retina. We performed this experiment either with no added lactate or with 1, 5, or 10 mM unlabeled lactate added to medium on the apical side (**Fig. 6A**). Consistent with our hypothesis, lactate added to the apical medium substantially enhances transport of ^13^C glucose from the basolateral to the apical side of the hfRPE cells (**Fig. 6B)**. Unlabeled lactate also suppresses the release of ^13^C lactate on the apical side (**Fig. 6C**).

**Fig. 6.**
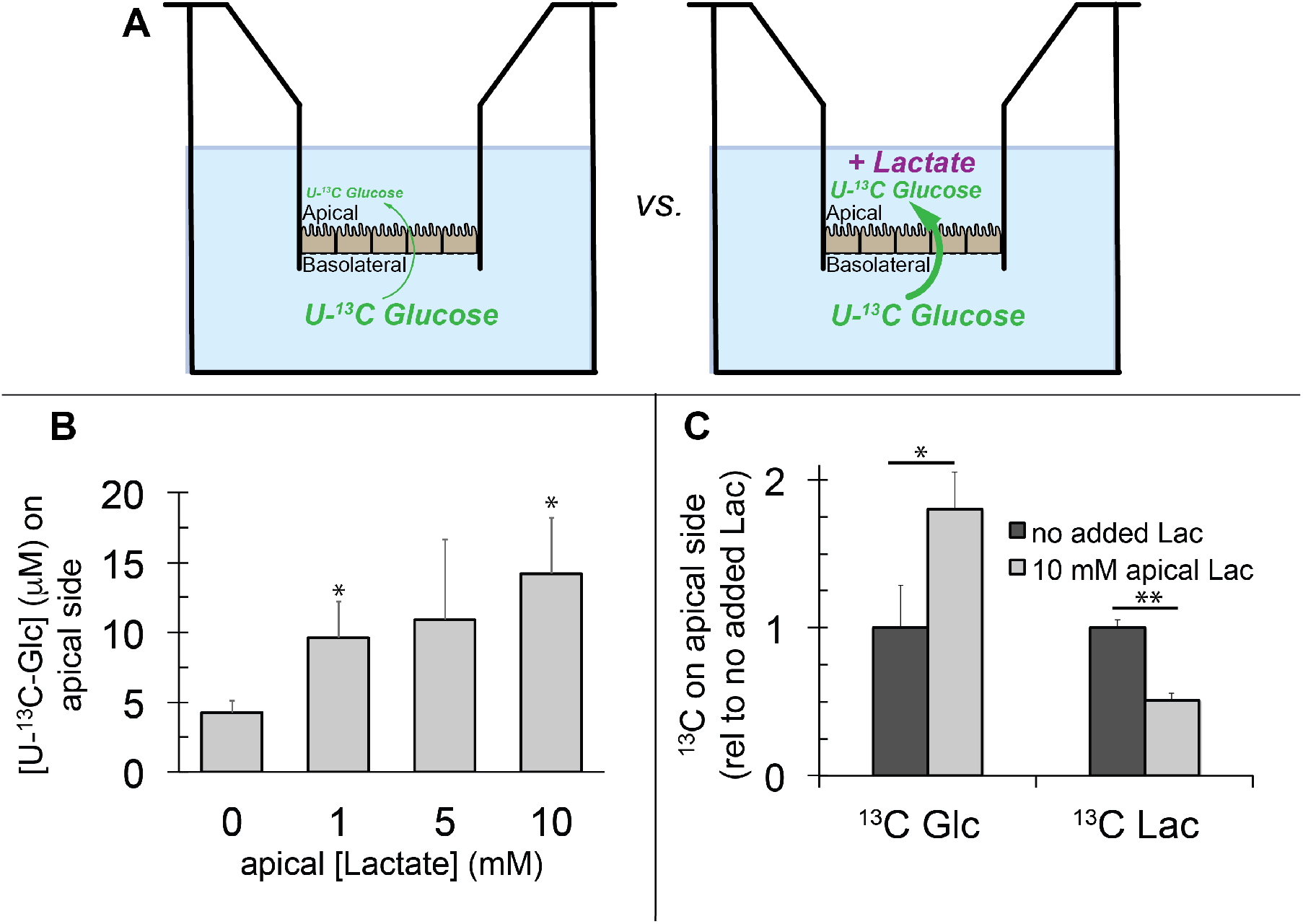
Lactate can enhance transport of glucose across a monolayer of RPE cells. **A**. Strategy to evaluate the effect of lactate on transport of glucose across a monolayer of RPE cells. Without lactate (left) glycolysis consumes glucose before it can cross the RPE cell monolayer. With lactate on the apical side (right) glycolysis is partially suppressed so more glucose can cross the monolayer without being consumed by glycolysis. **B.** Lactate (unlabeled) on the apical side enhances transport of U-^13^C glucose (U-^13^C Glc) (measured after 24 hours) from the basolateral to apical side of the RPE cell monolayer (n = 3 for each condition; representative of 3 separate experiments, error bars represent StDev * indicates p< 0.05 compared to no added lactate). **C.** Addition of lactate (unlabeled) to the apical side enhances transport of ^13^C Glc from the basolateral side to the apical side (left) (measured after 8 hours) and it suppresses the formation and/or release of ^13^C lactate (right) into the medium on the apical side. (n = 3 for each data point, error bars represent StDev).

### Long-term exposure to lactate enhances maturation of hfRPE cells

Culture conditions for hfRPE cells have been optimized for differentiation and for growth (35). High concentrations of lactate generally are not included in culture media for RPE cells. However, in their native environment within a living eye, RPE cells normally would be exposed to constantly high concentrations of lactate released from the retina.

Since lactate is metabolized efficiently by hfRPE cells we hypothesized that the presence of lactate in the culture medium might influence their differentiation state. To test this we incubated hfRPE cells in standard medium containing glucose until they were confluent and had achieved a trans-epithelial resistance ≥200 Ω•cm^2^. We then lowered the concentration of glucose to 0.5 mM and added 10 mM lactate. Remarkably, cell survival was excellent. After 21 days pigmentation of the lactate supplemented hfRPE cells increased substantially compared to control hfRPE cells provided with standard 5 mM glucose sans lactate. (**Fig 7A**). **Fig. 7B** shows that the levels of several key metabolic enzymes decreased, whereas the ratio of IDH3 to other IDH isoforms increased. Exposure to lactate also alters metabolic flux through decarboxylation and carboxylation of pyruvate (**Fig. 7C**) and reductive carboxylation (**Fig. 7D**). A more complete analysis of the effects of lactate on RPE cell differentiation is needed, but our initial findings, shown here, demonstrate that lactate not only supports RPE metabolism, but it also influences the differentiation state of RPE cells.

**Fig. 7.**
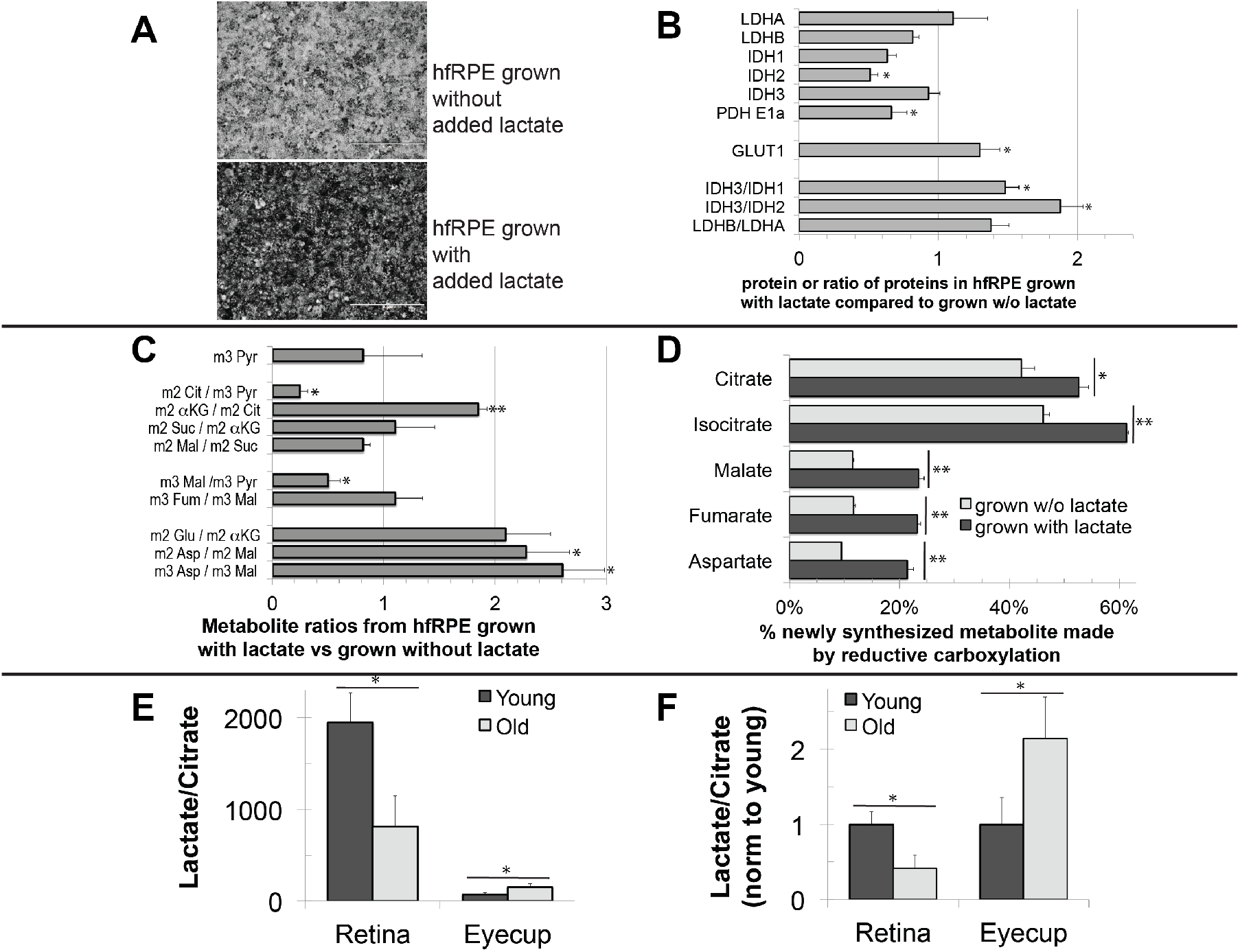
Effect of long-term exposure to lactate on hfRPE cells. A. hfRPE cells were grown on standard media to confluence and with a transepithelial resistance >200 O* cm^2^. They then were treated with either media containing 5 mM glucose (top) or 0.5 mM glucose/10 mM lactate (bottom) for 14 days. The pigmentation of the hfRPE cells treated with lactate increased substantially. **B.** Immunoblot quantification of protein extracted from hfRPE cells revealed that several key metabolic enzymes decreased in abundance with lactate supplementation (normalized to total protein) (n = 4; error bars represent SEM). The ratio of LDH3 to other LDH isoforms increased substantially. **C**. Quantification of metabolite ratios from hfRPE cell monolayers incubated with U-^13^C lactate for 10 min using GC-MS. The data show that lactate pretreatment alters metabolic flux through specific reactions. **D**. Lactate treatment enhances reductive carboxylation. The graph shows quantification of newly synthesized metabolites from hfRPE cell monolayers incubated with U-^13^C glutamine for 1 h using GC-MS. The percentage of newly synthesized metabolite made by reductive carboxylation was determined from m5 citrate and isocitrate or m3 malate, fumarate and aspartate relative to all other isotopomers of those metabolites that had incorporated ^13^C. **E**. Comparison of total Lactate/Citrate ratios in retina and eyecups from young (6 month) and old (32 month) mice after incubating the retinas or eyecups with 5 mM Glc and 2 mM Gln for 1 hour. **F**. Same data as in (E) normalized to the Lac/Cit ratio for young mice. (n = 2 for 6 month and n = 4 for 32 month; error bars represent StDev.)

### Metabolic specializations of the retina and RPE decline with age

The analyses of RPE metabolism in this study focused on the cultured hfRPE, which is a well characterized model that has been used to evaluate RPE metabolism (32). hfRPE cells have been used as a cell culture model for studying various diseases, including age-related macular degeneration (34). The RPE cultures used in the experiments reported here are of a similar age in culture as the ones used in other published studies, including those modeling AMD.

However, to confirm that the metabolic differences between retina and hfRPE are similar to the differences between retina and RPE in an eye, we used a mouse eyecup preparation with retinas removed to confirm the basic metabolic features that we identified in this study. We incubated freshly separated retinas and eyecups in medium containing glucose and glutamine and then analyzed metabolites by GC-MS. **Fig. 7E** compares the ratio of total lactate to total citrate in the retina vs. in the eyecup. Similar to the ratio we found for mouse retina and hfRPE, the lactate/citrate ratio is ~30 times higher in retina than in the eyecup.

Based on our findings we speculated that degeneration of photoreceptors in response to mutations or to aging could be influenced by the metabolic relationship between the retina and RPE. The eyecup preparation allowed us to examine the effect of aging on the lactate/citrate ratio in retina and RPE. **Figs. 7E and 7F** show that lactate/citrate in the retina *decreases* substantially as mice age from 6 months to 32 months. In contrast, lactate/citrate *increases* with age in the eyecup. In other words the retina becomes less glycolytic with aging while the eyecup, containing the RPE layer, becomes more glycolytic. These results suggest that the specialized metabolic features, that are fundamental to the retina-RPE ecosystem, decline with age both in the retina and in the RPE and that this decline could contribute to age-related vision loss (37, 38).

## DISCUSSION

### Model for a network of metabolic interdependence between the retina and RPE

**Fig. 8** summarizes our model for the retina-RPE metabolic ecosystem. Lactate from photoreceptors suppresses glycolysis in the RPE so more glucose can reach the retina.

**Fig. 8.**
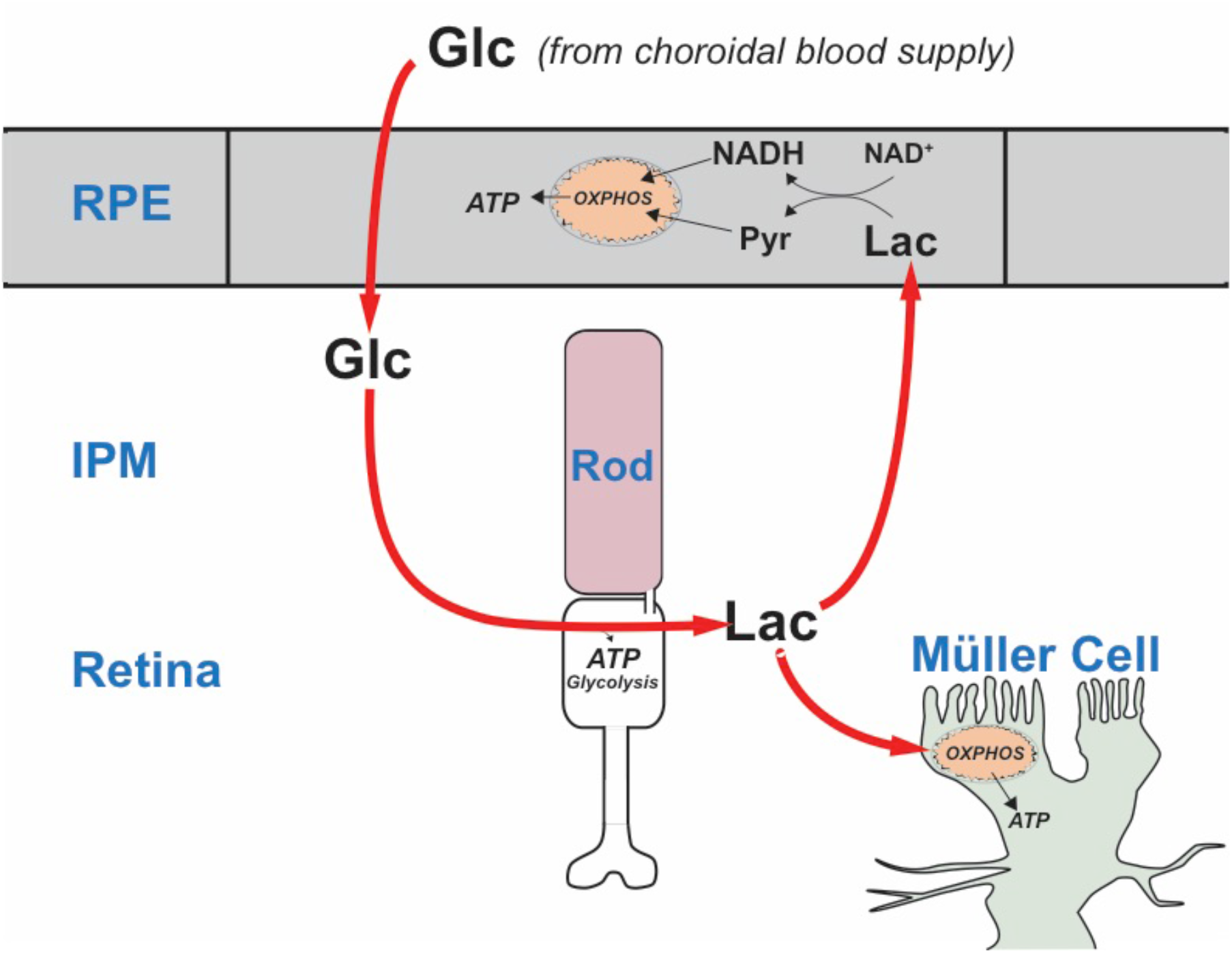
A working model that describes the flow of metabolic energy in the retina-RPE ecosystem. Photoreceptors convert glucose into lactate and release the lactate into the interphotoreceptor matrix. Lactate suppresses glycolysis in RPE cells by depleting NAD^+^. Lactate also fuels metabolic activity in Müller cells, which lack key enzymes that would be required for glycolysis.

### Previous evidence that cells in the retina have specific metabolic roles

The distributions of metabolic enzymes in mouse retina indicate that photoreceptors have the enzymes and transporters they need for glycolysis, but MGCs do not. Glycolysis requires pyruvate kinase (PK). The M2 isoform of PK (PKM2) is highly enriched in photoreceptors (10, 18–20, 39), but MGCs do not express substantial amounts of any PK isoform (10). MGCs also do not express hexokinase (20). Furthermore, lactate, rather than glucose, is the most effective source (10) of carbon for glutamine synthesis by MGCs (40) in mouse retinas. Based on these observations, we proposed that MGCs in a retina primarily use lactate produced by photoreceptors (41). Together with the results described in this report it appears that the central role of photoreceptors in retinal energy metabolism is to convert glucose to lactate, which then is used as fuel by both RPE and Müller cells.

### Significance of aerobic glycolysis in the retina

Enhanced capacity for anabolic metabolism has been proposed as the purpose of aerobic glycolysis in photoreceptors (10, 18, 19), but our model suggests an additional purpose. We propose that the laminated structure of the eye, in which the RPE separates the retina from its source of nutrients, requires photoreceptors to produce and release lactate to suppress glycolysis so that sufficient glucose can flow through the RPE.

### The “retinal ecosystem” model helps to explain recent findings from genetic manipulations of photoreceptors and RPE

Engineering photoreceptors to be more glycolytic makes them more robust, whereas engineering RPE cells to be more glycolytic causes photoreceptors to degenerate (3, 21–23). The model shown in **Fig. 8** provides an explanation for those seemingly opposing findings. When photoreceptors are engineered to be more glycolytic they produce more lactate that can more substantially suppress glycolysis in the RPE.In this way more glucose reaches the retina, thereby enhancing photoreceptor survival. When the RPE is more glycolytic it consumes more glucose, leaving less glucose available for the retina so photoreceptors starve and become stressed.

### The concept of a metabolic ecosystem and its relationship to retinal disease

The “retina ecosystem” model in **Fig. 8** suggests an explanation for the linkage between Age-Related Macular Degeneration and accumulation of mitochondrial DNA damage in RPE cells (42). Photoreceptors may starve when RPE mitochondria fail because the RPE becomes more dependent on glycolysis, which prevents glucose from reaching the retina.

The concept of a metabolic ecosystem also has implications for other types of retinal disease. Mutations that affect genes expressed only in rods can cause rods to degenerate. However, cones subsequently degenerate as a consequence of the loss of rods, even though the cones are not affected directly by the mutant gene (43). One reason for this is that loss of a cone viability factor that normally is produced by rods may contribute to cone degeneration in this type of disease state (44). The model in **Fig. 8** suggests another factor that also can contribute to the secondary loss of cones when rods degenerate. A retina without rods makes less lactate (8). We have shown in this report that, without lactate to suppress glycolysis, RPE cells oxidize more glucose and transport less. In a rod-less retina the loss of lactate production would limit the rate at which glucose can be supplied to cones. This can explain why cones starve (2) and why 2-NBDG accumulates in RPE cells (45) when rods degenerate. Consistent with this explanation, cones in these retinas can be rescued from degeneration by supplementation with an alternative supply of glucose (45).

### Other fuels also may contribute in the metabolic ecosystem

This study highlights one way that RPE, photoreceptors and MGCs can work together as an ecosystem of metabolically specialized interdependent cells. Our investigation focused on glucose and lactate, but it is very likely that fatty acids, ketone bodies, (46–48) and metabolites from other metabolic pathways (20) also contribute significantly to this metabolic ecosystem. For example, oxidation of fatty acids by the RPE can supply the retina with ketone bodies (46). Furthermore, a recent study showed that RPE oxidizes fatty acids from photoreceptor phagocytosis, (48) which also can suppress the need for RPE cells to consume glucose. Together these studies suggest that energy homeostasis in retina and RPE relies on a specialized metabolic interplay that could form the basis of future therapies for a range of retinal degenerative diseases.

## MATERIAL AND METHODS

### Animals

All research was authorized by the University of Washington Institutional Animal Care and Use Committee. Mice in the C57BL6 background were maintained in the University of Washington South Lake Union vivarium at 27.5 °C on a 14h/10h light-dark cycle. Transgenic mice expressing eGFP under the Nrl promoter (49) (RRID:IMSR_JAX:021232), or tdTomato under the Rlbp-CRE promoter (28) were described previously. Aged mice (6 month and 32 month) were C57BL/6J. This strain does not carry the rd8 mutation in the Crb1 gene (50).

Transgenic heterozygote zebrafish in the AB background were maintained in the University of Washington South Lake Union aquatics facility at 27.5 °C on a 14h/10h light-dark cycle. Fish used for experiments were male and female siblings between 12-24 months old. A transgenic line stably expressing tdTomato in Müller cells (GFAP:tdTomato; RRID:ZDB-FISH-150901-17843) was described previously (30). Prior to gavage experiments, fish were fasted >18 h and dark-adapted >12 h.

### Antibodies

Arrestin1, D9F2 (from Larry Donoso and Cheryl Craft) IHC: 1:200
GLUT1, (AbCam, ab115730; RRID:AB_10903230) IB: 1:200,000, 0.86 ng/ml; IHC 1:1000, 0.17 mg/ml
GLUT3, (AbCam, ab41525; RRID:AB_732609) IB: 1:5000, 0.136 ug/ml
GLUT4, (AbCam, ab654; RRID:AB_305554) IB: 1:5000
Glutamine synthetase, (Millipore, MAB302; RRID:AB_2110656) IHC: 1:1000
MTCO1 (Abcam, ab14705; RRID:AB_2084810)

## Tissue preparations for immunoblotting

Frozen tissue samples were homogenized in RIPA buffer (150 mM NaCl, 1% Triton X-100, 0.05% sodium deoxycholate, 0.1% SDS, 50 mM Tris, pH 8.0 with a mixed phosphatase/protease inhibitor cocktail (ThermoFisher 88668), briefly sonicated, then rocked at 4°C for 30min. Samples were then spun at 13,300 RPM at 4°C for 15 min, and the supernatant was normalized for loading by BCA assay to 20 mg/tissue. RPE protein lysate was prepared according to a described protocol (51).

To prepare membrane fractions, frozen tissue samples were homogenized in PBS (0.14 M, pH 7.4) with a mixed phosphatase/protease inhibitor cocktail, then rocked at 4°C for 30 min. Samples were then spun at 45,000 rpm at 4°C, the supernatant (cytosolic fraction) drawn off and saved, and the pellet (membrane fraction) was resuspended in an equal volume of PBS. After mixing with 5X Laemmli loading buffer, 1 ml benzonase (Millipore 70746) was added. Each tissue was then loaded with equal volumes of cytosolic and membrane fraction.

## Immunoblotting

Samples were run on 12%, self-cast acrylamide gels and transferred onto PVDF membranes (Millipore IPFL00010). Following protein transfer, membranes were blocked with LI-COR Odyssey Blocking Buffer (LI-COR, 927-40000) for 1 h at room temperature. Primary antibodies were diluted in blocking buffer and incubated overnight at 4°C. Membranes were washed, incubated with secondary antibody (LI-COR IRDye 800CW, 926-32210, (RRID:AB_621842), and 926-32211, (RRID:AB_621843),1:5000 1h at room temperature, and washed again. Imaging was performed using the LI-COR Odyssey CLx Imaging System (RRID:SCR_014579).

## Immunohistochemistry

Retinal eyecups were micro-dissected from C57BL6J mice and were fixed in 4% paraformaldehyde in PBS, rinsed with PBS, incubated in a sucrose gradient (5%, 10%, and 20%), embedded into OCT and cryosectioned at 20pm. Mouse sections were washed in PBS, then blocked in IHC buffer (5% normal donkey serum diluted in PBS with 2 mg/mL BSA and 0.3% Triton X-100) for 1 h. Primary antibodies were diluted in IHC blocking buffer as specified, and applied to blocked cryosections overnight at 4°C. Secondary antibodies were diluted at 1:3000 in IHC blocking buffer, and applied to mouse retina sections for 1 h in darkness. Sections were washed in PBS three times, and mounted with SouthernBiotech Fluoromount-G (Fisher Scientific) under glass coverslips and visualized using a Leica SP8 confocal microscope with a 63X oil objective. Images were acquired at a 4096x4096 pixel resolution with a 12-bit depth using Leica LAS-X software (RRID:SCR 013673).

## RPE cell culture

Human fetal eyes with a gestational age of 16-20 weeks were harvested and shipped overnight on ice in RPMI media containing antibiotics from Advanced Bioscience Resources Inc. (Alameda, CA).Dissections of fetal tissue were performed within 24 hours of procurement and followed a modified version of the dissection protocol in order to isolate the retinal pigment epithelium (RPE) (35). The fetal RPE sheets were incubated at 37°C with 5% CO_2_ and cultured in RPE media. The RPE media consisted of Minimum Essential Medium alpha (Life Technologies) supplemented with 5% (vol/vol) fetal bovine serum (Atlanta Biologicals), N1-Supplement (Sigma-Aldrich), Nonessential Amino Acids (Gibco), and a Penicillin-Streptomycin solution (Gibco). Isolated fetal RPE reached confluency about 3-4 weeks after dissection and was then passaged using a 0.25% Trypsin-EDTA solution (Gibco) and passed through a 40 mm nylon cell strainer (BD Falcon) in order to collect a suspension of single cells. After counting, the RPE cells were plated onto 0.3 cm^2^ cell culture inserts (Falcon) coated with Matrigel (Corning) at a seeding density of 100,000 cells per insert. Cells grown on these inserts were cultured in RPE media containing 1% (vol/vol) FBS. Transepithelial resistance was measured weekly after 2 weeks in culture using a Millicell ERS-2 Epithelial Volt-Ohm Meter (Millipore).

## Oral Gavage

Mice were fasted overnight in the dark, and gavaged the next morning in ambient light. A micro-syringe fitted with a 22 gauge 1.5” straight 1.25 mm ball-tip needle was used to orally administer 100 μl of 50 mM 2-NBDG (Invitrogen) dissolved in water. Successfully gavaged mice were returned to darkness during the 2-NBDG incubation period.

Zebrafish were gavaged using methods described previously(52) under red light. Briefly, overnight fasted adult zebrafish were anaesthetized > 1 min with 150 mg/mL MS-222 in fish water. Fish were placed in a slit cut in a cellulose sponge soaked with MS-222 solution, and the sponge was rotated to orient the fish mouth up. A micro-syringe fitted with thin, flexible 1 mm OD plastic tubing was used to orally administer 5 μL of either fish water or 30 mM 2-NBDG (Invitrogen). Gavaged fish were immediately placed into a recovery tank of fresh fish water and monitored briefly using a UV flashlight for regurgitation of 2-NBDG. Successfully gavaged fish were returned to darkness during the 2-NBDG incubation period.

## Tissue slicing and imaging

Gavaged mice were euthanized by asphyxiation with CO_2_. Zebrafish were euthanized in an ice bath followed by cervical dislocation. Euthanized animals were enucleated, and the retinas dissected away under red light into cold Ringer’s solution (133 mM NaCl, 2.5 mM KCl, 1.5 mM NaH_2_PO_4_, 2 mM CaCl_2_, 1.5 mM MgCl_2_, 10 mM HEPES, 10 mM D-glucose, 1 mM sodium lactate, 0.5 mM L-glutamine, 0.5 mM reduced glutathione, 0.5 mM sodium pyruvate, 0.3 mM sodium ascorbate, pH 7.4). Isolated retinas were mounted on filter paper (0.45 μm pore, mixed cellulose, Millipore) and flattened with gentle suction. After peeling away remaining RPE, flat-mounted retinas were sliced into 300-400 μm slices using a tissue slicer (Stoelting). Slices were rotated 90**°** and the filter paper edges buried in strips of wax on a coverslip for imaging at room temperature. Fresh retinal slices were imaged at room temperature using a Leica SP8 confocal microscope with a 40X water objective; excitation/emission wavelengths were 488/525-575 nm for 2-NBDG, and 559/580-630 nm for tdTomato. Leica LAS-X (RRID:SCR_013673) software was used to acquire images at 2048 x 2048 pixel resolution with 12 bit depth, and Z-stacks imaged every 0.5 μm over a tissue depth of 10-30 μm.

## Image analysis

ImageJ software (RRID:SCR_002285) was used for quantification of 2-NBDG fluorescence in fresh retinal slices. 10 slices of each Z-stack were maximum intensity projected, and retinal layers were identified by morphology and expression of transgenic markers. For every slice, 3 small uniformly sized rectangular regions of interest (ROIs) were placed randomly in each retinal layer, and mean fluorescence intensity of each ROI was measured. Average 2-NBDG fluorescence in each layer was divided by the autofluorescence of corresponding retinal layers from animals gavaged with saline or water.

## Metabolic flux analysis

Isolated mouse retina or confluent human fetal RPE cells were changed into pre-warmed Krebs-Ringer Bicarbonate Buffer containing, depending on the experiment, [1,2] ^13^C glucose, U-^13^C glucose, U-^13^C lactate or U-^13^C glutamine (Sigma) as described elsewhere (5, 7, 9). Both retinas and RPE cells were incubated for 5 min, 30 min, 60 min and 120 min. Metabolites from each time point were extracted and analyzed by gas chromatography mass spectrometry (GC-MS, Agilent 7890/5975C) as described in detail (5, 6).

## Measurement of U-^13^C glucose transport across hfRPE cells on transwell filters

After maturation for 4-6 weeks in culture, hfRPE cells grown on transwell filters were changed into 500 ml of DMEM containing 1% FBS on each side. 5.5 mM U-^13^C glucose (Cambridge Isotope Laboratories) was included in the medium in the basolateral side while various concentrations of sodium lactate was added to the apical side, while maintaining a constant pH. Apical side medium was collected at 8 and 24 h to analyze the transported U-^13^C glucose by liquid chromatography coupled with triple quadrupole mass spectrometry (Waters Xevo TQ Tandem mass spectrometer with a Waters ACQUITY system with UPLC) as reported in detail (7).

## Serial Block Face Scanning SEM

Mouse eyes were enucleated, the anterior half was dissected away, and the eyecup was cut in half. Tissue was fixed in 4% glutaraldehyde in 0.1 M sodium cacodylate buffer, pH 7.2, at room temperature (RT), then stored overnight at 4°C. Samples were washed 4 times in sodium cacodylate buffer, postfixed in osmium ferrocyanide (2% osmium tetroxide/3% potassium ferrocyanide in buffer) for 1 h on ice, washed, incubated in 1% thiocarbohydrazide for 20 min, and washed again. After incubation in 2% osmium tetroxide for 30 min at RT, samples were washed and en bloc stained with 1% aqueous uranyl acetate overnight at 4°C. Samples were finally washed and en bloc stained with Walton’s lead aspartate for 30 min at 60°C, dehydrated in a graded eth-anol series, and embedded in Durcupan resin. Serial sections were cut at 60 nm thickness and imaged with 6 nm pixel size using a Zeiss Sigma VP scanning electron microscope fitted with a Gatan 3View2XP ultramicrotome apparatus. Imaged stacks were concatenated and aligned using TrakEM2 (RRID:SCR_008954). Unless stated otherwise, five washes with water were used for all wash steps.

## Statistical analyses

R (RRID:SCR_001905) with R Commander was used to perform one-way ANOVA for NBDG gavage experiments.

### Reproducibility

All data shown here have been reproduced at least three times by the authors.

### Data availability

All data supporting the findings of this study are available within the paper.

## Acknowledgements

This study was supported by funding from NIH EY06641 and NIH EY017863 to JBH, NIH EY026020 to SEB and from NEI core grant EY001730.

